# Discovery of novel brain permeable human ACSS2 inhibitors for blocking breast cancer brain metastatic growth

**DOI:** 10.1101/2023.12.22.573073

**Authors:** Emily Esquea, Lorela Ciraku, Riley G. Young, Jessica Merzy, Alexandra N. Talarico, Adel Ahmed Rashad, Simon Cocklin, Nicole L. Simone, Joris Beld, Mauricio J. Reginato, Alexej Dick

## Abstract

Breast-cancer brain metastasis (BCBM) poses a significant clinical challenge, resulting in an end-stage diagnosis and hindered by limited therapeutic options. The blood-brain barrier (BBB) acts as an anatomical and physiological hurdle for therapeutic compounds, restricting the effective delivery of therapies to the brain. In order to grow and survive in a nutrient-poor environment, tumors in the brain must adapt to their metabolic needs, becoming highly dependent on acetate. These tumors rely on the conversion of acetate to acetyl-CoA by the enzyme Acetyl-CoA synthetase 2 (ACSS2), a key metabolic enzyme involved in regulating fatty acid synthesis and protein acetylation in tumor cells. ACSS2 has emerged as a crucial enzyme required for the growth of tumors in the brain. Here, we utilized a computational pipeline, combining pharmacophore-based shape screen methodology with ADME property predictions to identify novel brain-permeable ACSS2 inhibitors. From a small molecule library, this approach identified 30 potential ACSS2 binders, from which two candidates, AD-5584 and AD-8007, were validated for their binding affinity, predicted metabolic stability, and, notably, their ability to traverse the BBB. We show that treatment of BCBM cells, MDA-MB-231BR, with AD-5584 and AD-8007 leads to a significant reduction in lipid storage, reduction in colony formation, and increase in cell death *in vitro*. Utilizing an *ex vivo* orthotopic brain-slice tumor model, we show that treatment with AD-8007 and AD-5584 significantly reduces tumor size and synergizes with radiation in blocking BCBM tumor growth *ex vivo.* Importantly, we show that following intraperitoneal injections with AD-5584 and AD-8007, we can detect these compounds in the brain, confirming their BBB permeability. Thus, we have identified and validated novel ACSS2 inhibitor candidates for further drug development and optimization as agents for treating patients with breast cancer brain metastasis.

## Introduction

Breast cancer is the most commonly diagnosed cancer in women worldwide [1], with an estimated 10-35% of breast cancer patients developing metastasis to the brain [2–4]. Breast-cancer brain metastases (BCBM) currently represent an incurable event [5], with over 80% of patients losing their lives within a year of diagnosis [1, 6]. Limited therapeutic interventions, including surgical resection, whole brain radiation, and chemotherapy, are often ineffective and yield detrimental effects on healthy brain tissue, thereby profoundly diminishing the quality of life of affected patients [5, 7, 8]. Thus, there is an urgent need to identify small molecules that can block tumor growth in the brain.

Brain tumor cells are situated within a nutrient-depleted and hypoxic tumor brain microenvironment; thus, these cancer cells must adapt metabolically to survive [9, 10]. Metabolic reprogramming in cancer cells in the brain allows for the survival, growth, and progression of these tumors but also leads to metabolic vulnerabilities that may be exploited therapeutically [9, 10]. One of these metabolic vulnerabilities arises from the acetate dependency of brain tumor cells [11, 12]. Acetate is an alternative carbon source to generate acetyl-CoA via the nuclear-cytosolic enzyme acetyl-CoA synthetase 2 (ACSS2)[11–13]. Acetyl-CoA plays a critical role in cellular energetics, leading to the generation of *de novo* lipid and fatty acids, and can be utilized in protein and histone acetylation [13, 14]. ACSS2 has been shown to play a critical role in tumor cell growth in various cancers, including breast [15–17] and brain cancers [18]. In particular, ACSS2 may serve as an attractive therapeutic target for tumors in the brain, such as glioblastoma and brain metastasic tumors, due to the preferential use of acetate as an energy source in these tumors [19]. Importantly, genetically targeting ACSS2 in brain tumors has previously been shown to block tumorigenesis [18, 20]. Additionally, ACSS2 null mice are phenotypically normal, without embryonical or developmental deficiencies, suggesting ACSS2 may be a nonessential gene under normal conditions [21], thus making ACSS2 an attractive cancer-specific target. Several small molecules targeting ACSS2 have been identified and tested in liver [15] and breast cancer models [16, 17]. Currently, one ACSS2 inhibitor, MTB-9655, the first oral ACSS2 inhibitor, is in phase one clinical trial for advanced solid tumors [22]. However, to our knowledge, there have not been small molecule ACSS2 inhibitors that can cross the blood-brain barrier previously identified for treating cancers in the brain.

In this study, we sought to discover novel, pharmacologically stable, small-molecule inhibitors of ACSS2 that are able to cross the blood-brain barrier. Using a previously validated computational workflow [23–27] with the addition of brain and CNS-specific parameters, we identified several new chemotypes in the ACSS2 inhibitor class. These ACSS2 inhibitor analogs exhibit drug-like properties in line with computational predictions and have been validated in several BCBM models. We show that these inhibitors can suppress BCBM tumor growth *in vitro* and *ex vivo* and cross the blood-brain barrier *in vivo*. These first-in-class BBB-permeable ACSS2 inhibitor analogs provide scaffolds for further optimization of ACSS2-targeting chemotypes and enable improved cancer treatments in the brain.

## Results

### Formulating a Computational Pipeline and Predicting Drug-like Properties for the Discovery of Brain-Permeable ACSS2 Inhibitors

The quinoxaline-based chemotype VY-3-249 targeting ACSS2 previously identified [15] is not predicted to traverse the blood-brain barrier as it has low oral CNS scoring profiles (**Fig. 1A and Fig. S1**). Although, a new derivative of VY-3-249, compound VY-3-135, a quinoxaline-based chemotype was recently shown to have increased potency and stability compared to VY-3-249 [28], yet it is also not predicted to cross the blood-brain barrier (**Fig. 1A and Fig. S1**). Thus, we sought to identify novel ACSS2 inhibitor chemotypes with blood-brain barrier permeability in order to target brain tumors. We utilized a pharmacophore-based shape screen methodology [29–31], further processed through validation of binding poses and computational prediction of [30] absorption, distribution, metabolism, and excretion (ADME) properties and other drug-like features (**Fig. 1B**). These *in silico* predictions were conducted employing StarDrop V7 (Optibrium, Ltd., Cambridge, UK), with the incorporation of the oral central nervous system (CNS) drug profile and an auxiliary parameter for logD [23, 32, 33]. The oral CNS drug profile comprises numerous models integrated into an overall score by a probabilistic scoring algorithm. This scoring system spans from 0 to 1, where a score of 0 suggests a non-drug-like compound, while 1 indicates the paradigm of a drug. This computational pipeline and stringent drug-like properties filtering distilled our initial molecule pool to 30 potential ACSS2 binders.

**Figure 1:**
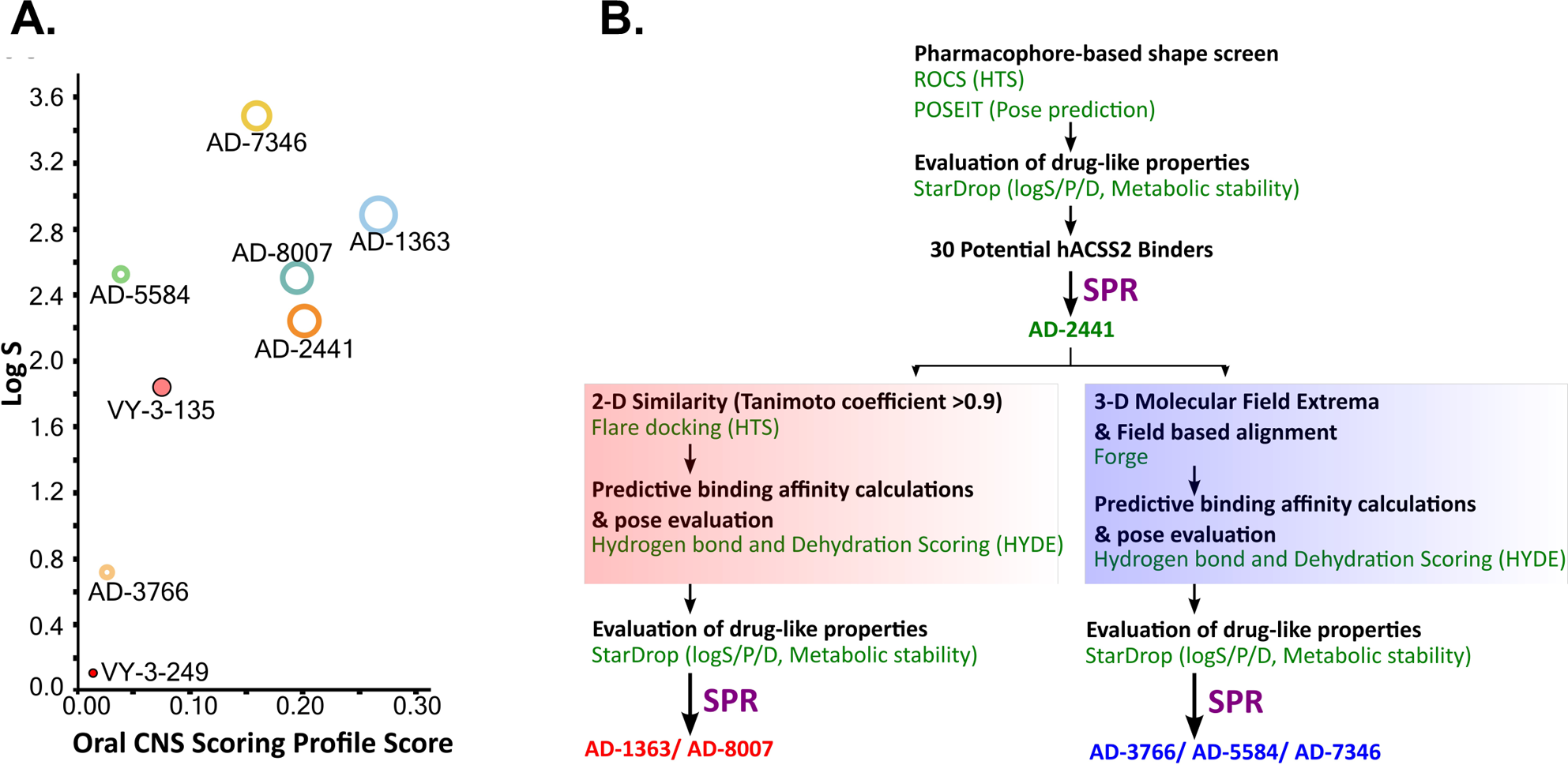
Computational pipeline and predicted drug-like properties for the discovery of AD-2441 and its analogs. **(A)** Oral CNS Scoring profile score vs. logS of AD-2441 and its analogs, including two currently known control inhibitors, VY-3-249 and VY-3-135. Plot showing the StarDrop V7 (Optibrium, Ltd., Cambridge, UK)–derived logS versus a multimetric oral CNS profile score. Score composition and importance of each contributor: logS = 0.8, logP = 0.6, logD = 0.6, BBB category = 0.55, BBB log(brain:blood) = 0.55, P-gp category = 0.5, HIA = 0.4, hERG pIC50 = 0.2, 2D6 affinity category = 0.16, 2C9 pKi = 0.16, PPB90 category = 0.1. The size of the circle correlates with the corresponding score or probability. **(B)** Computational and SPR-based validation pipeline for the discovery of novel ACSS2 inhibitors.

### Validating the Binding Affinity and Predicted Metabolic Stability of ACSS2 Inhibitors

From the list of 30 potential ACSS2 inhibitors from our computational workflow (**Fig. 1B**), we next tested whether these compounds bind to their intended target, human ACSS2, in order to inhibit its function. To test the binding affinity of the potential ACSS2 inhibitors, we used surface plasmon resonance (SPR) interaction analysis for directing binding affinity and kinetic and a fluorescent-based ATPase inhibition assay to ascertain *in vitro* ATPase inhibition (IC_50_) (BellBrook Labs). The outcome identified six candidates exhibiting low-micromolar affinities and IC_50s_ in the high nanomolar range (**Fig. 2, 3**, and **Table S1**, note that due to suboptimal drug-like properties, AD-3766 was not used for downstream assessments).

**Figure 2:**
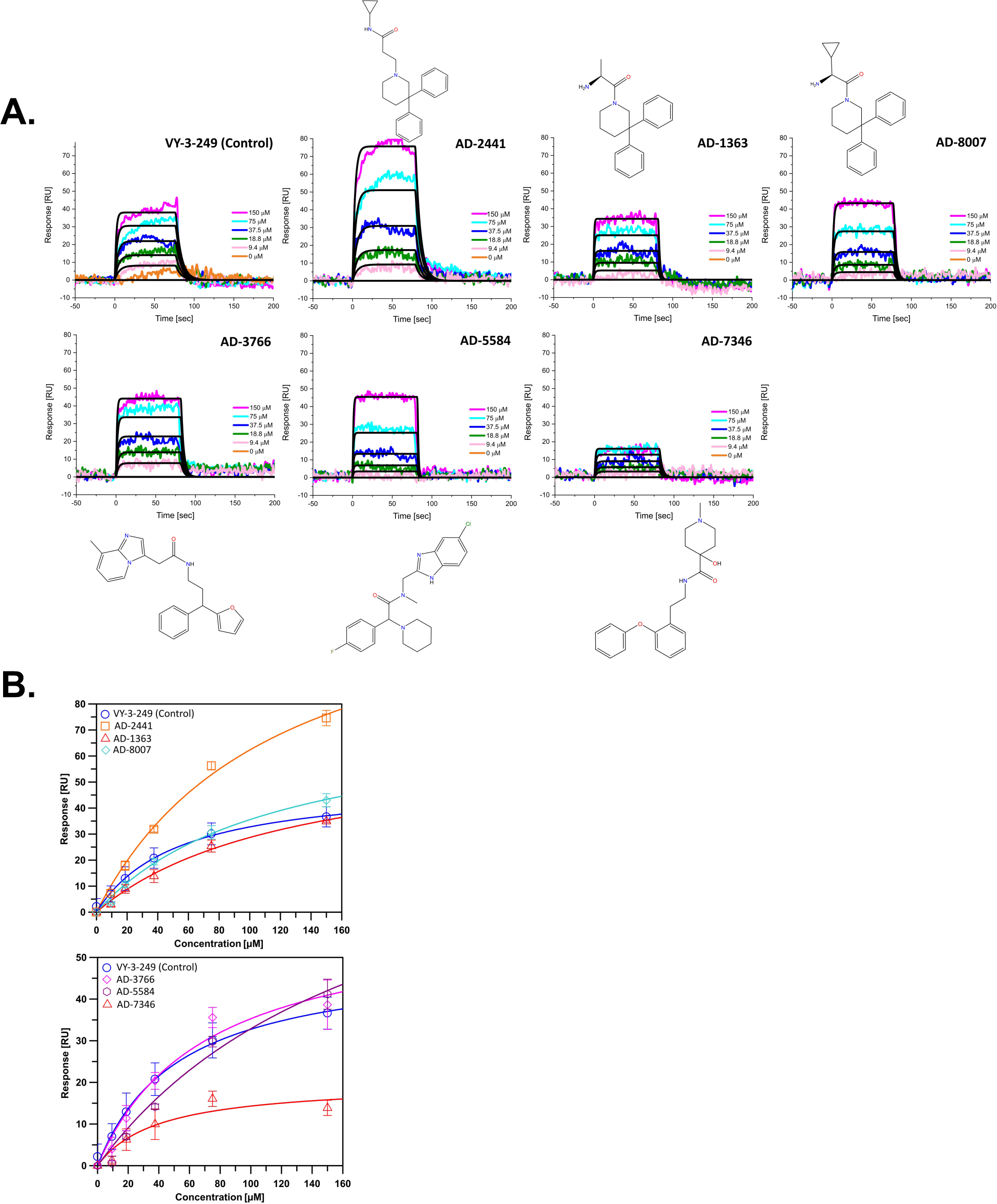
Representative sensorgrams and structures for AD-2441 and its analogs binding to ACSS2. (A) Sensorgrams of ACSS2 binders. Colored lines represent collected data from the dilution series, whereas black lines represent a fit to a 1:1 binding model—interaction parameters derived from a triplicate (n=3) of data given in Table S1. (**B**) Binding isotherms of AD-2441 and analogs. Binding isotherms are derived from panel A. Experiments were performed in triplicate, and data displayed with standard deviations (SD with n=3).

**Figure 3:**
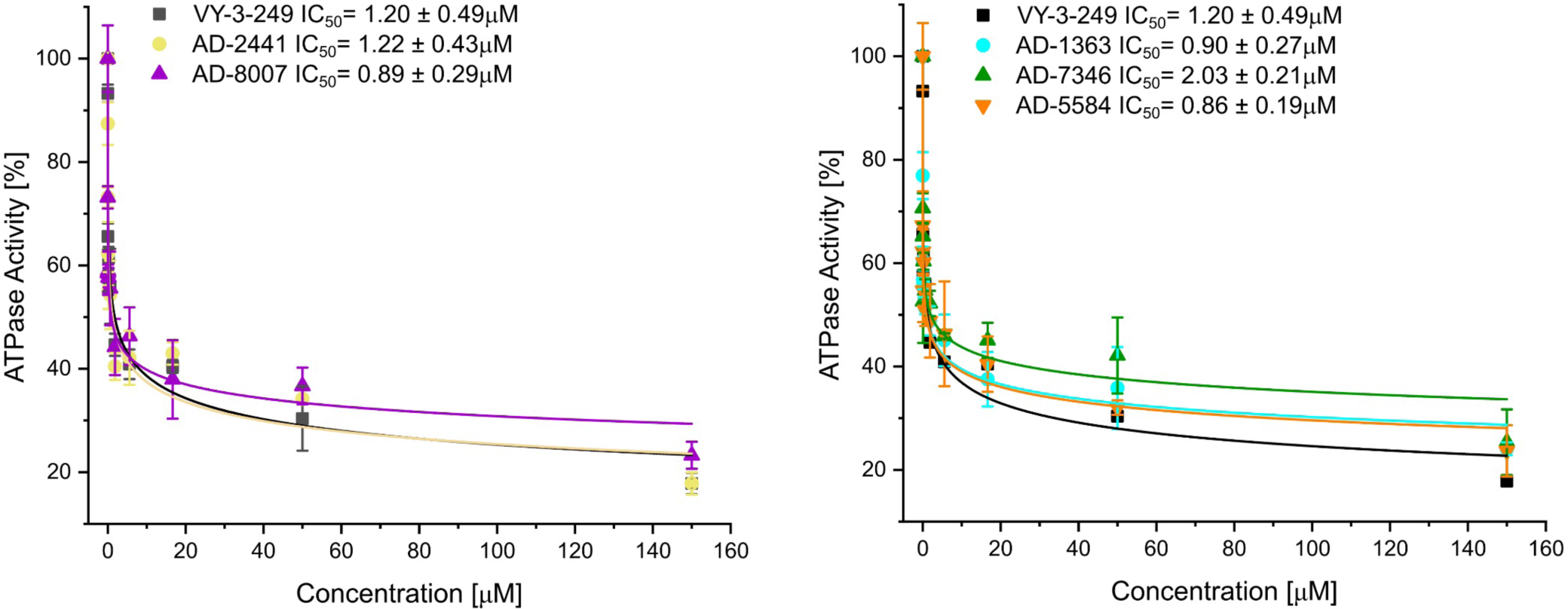
Fluorescent polarisation-based assay for measurement of inhibition of ATP to AMP conversion of ACSS2 by selected ACSS2 inhibitors. Experiments were performed using the Tecan Spark multimode microplate reader with 100 nM ACSS2 and varying compound concentrations. Experiments were performed in triplicate, and data displayed with standard deviations (SD with n=3).

ACSS2 belongs to an enzyme family known for initiating reactions that generate AMP through a two-phase process [34]. Initially, acyl-AMP is formed while simultaneously releasing pyrophosphate. The intermediate stage involves the formation of acetyl-AMP. Subsequently, Coenzyme A (CoA) replaces AMP, producing the endproduct acetyl-CoA [34]. ACSS2 comprises a C-terminal and N-terminal lobe with CoA and acetyl-AMP binding between those two domains (**Fig. 4A**). For our docking approach, we compared first the crystal structure of Adenosine-5’-propylphosphate (from PDB: 1PG4) with our docked Adenosine-5’-propylphosphate pose, to validate the quality of the docking approach (**Fig. 4B**) and highlighting the accuracy of our homology model of ACSS2 for compound docking.

**Figure 4:**
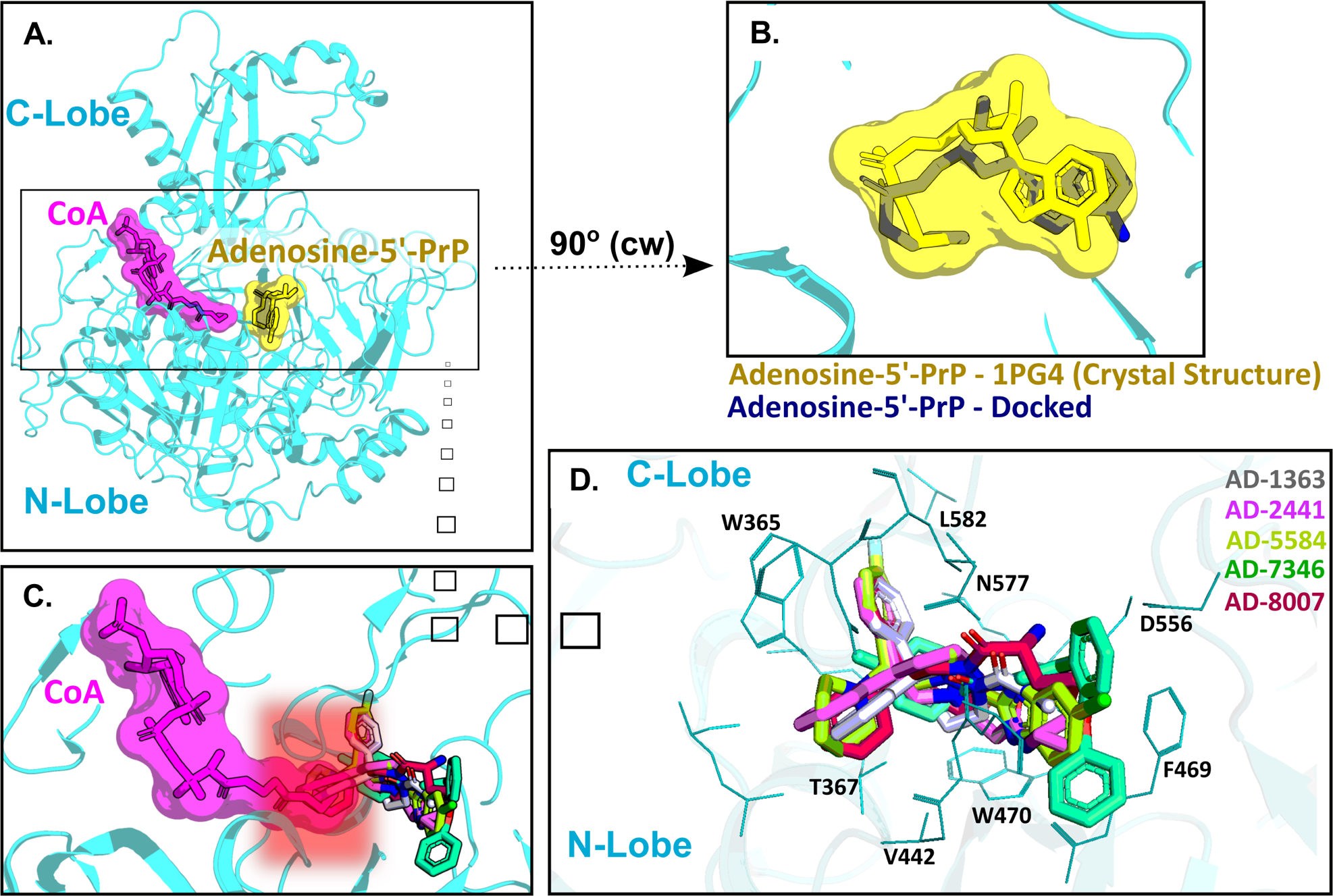
AD-2441 and its analogs are predicted to bind ACSS2 within the nucleotide-binding pocket and stabilized by various hydrophobic and polar contacts and aromatic stacking interactions. Docking calculations were performed into a homology model (using swissmodel.expasy.org)[50, 51] of ACSS2 based on the crystal structure of Salmonella enterica acetyl-CoA synthetase in complex with cAMP and Coenzyme A (PDB: 5JRH). (**A**) Homology model of ACSS2 with superimposed CoA (purple) and Adenosine-5’-propylphosphate (yellow, extracted from PDB: 1PG4). (**B**) Superimposed crystal structure of Adenosine-5’-propylphosphate (yellow, extracted from PDB: 1PG4) and docked (DiffDock and Flare version 5 minimized) Adenosine-5’-propylphosphate pose (blue). (**C**) All docked compounds are predicted to bind within the Adenosine-5’-propylphosphate site and additionally sterically interfere with CoA (purple) binding indicated by the red helo. (**D**) Close-up view of binding poses of AD-1363, AD-2441, AD-5584, AD-7346, and AD-8007 between the C- and N-Lobe of ACSS2.

Subsequent docking calculations predict that all six analogs bind near the acetyl-AMP site (**Fig. 4C**), potentially mimicking a short-lived transition state vital for ACSS2 function [35]. In addition, the binding of our compounds to ACSS2 most likely interferes sterically with CoA binding and, therefore, acetyl transfer (highlighted by the red helo). Besides hydrophobic (Val442, Leu582) and polar interactions (Asn577, Asp556), such as hydrogen bonds and salt bridges, a notable key feature of all compounds were π-π interactions via their bifurcated aromatic moieties with Trp365, Phe469, and Trp470 (**Fig. 4D**). Importantly, these potential inhibitors showcased predicted improved drug-like characteristics to those of control compounds with most notable metabolic stability (**Fig. S2A-C**) and BBB permeability (**Fig. 1A and S1C**).

Orally administered drugs undergo first-pass effect, a common occurrence in which drugs become metabolized, typically in the liver, and the concentration of the active drug is reduced [36]. Due to the first-pass effect, the metabolic stability of compounds can limit the concentration of these compounds in the bloodstream, directly affecting drug efficacy. Thus, we sought to computationally investigate if our identified ACSS2 inhibitors could potentially outperform previously established ACSS2 inhibitors, VY-3-249 and VY-3-135, in terms of predicted metabolic stability. We utilized a computational analysis, applying the P450 module in StarDrop V7 software (Optibrium, Ltd., Cambridge, UK) to predict each compound’s primary metabolizing Cytochrome P450 isoforms using the WhichP450™ model [24, 31, 33]. This was followed by an estimation of the compound’s affinity to that isoform through the application of the HYDE function in SeeSAR (BioSolveIT Gmbh, Sankt Augustin, Germany) [37](**Fig. S2**). This methodology has been used successfully in predicting and improving the metabolic stability of HIV-1 inhibitory compounds [23–27]. The CYP3A4/2D6 isoforms act as the major metabolizing enzyme for all compounds, including the control (**Fig. S2A**). We investigated the predicted metabolic lability of our compounds with the CY3A4 isoform, gauging the overall composite site lability (CSL) score and the number of labile sites. The CSL score amalgamates the labilities of individual sites within the compound, providing insight into the efficiency of the molecule’s metabolism [24, 31, 33]. Compared to the control VY-3-249, our compounds displayed lower labile sites and CSL scores, suggesting increased metabolic stability (**Fig. S2B, S2C**). While the CSL score and number of labile sites provide useful information, they assume all compounds bind with similar affinity to the CYP3A4 isoform. However, other factors influence metabolic stability, such as the binding affinity to the CYP3A4 isoform, compound reduction rate, and inherent compound properties like size and lipophilicity [24, 31, 33].

Consequently, we performed predictive binding affinity calculations using the hydrogen bond and dehydration (HYDE) energy scoring function in SeeSAR 12.1 (BioSolveIT Gmbh, Sankt Augustin, Germany)[37] with the structure of the human CYPA4 bound to an inhibitor (PDB: 4D78)[38]. The HYDE scoring function in SeeSAR offers a range of affinities, stipulating an upper and lower limit. By integrating the CSL scores, labile sites, and predicted CYP3A4/2D6 affinity, our analysis suggests that compounds AD-1363, AD-2441, AD-5584, and AD-8007 may have improved metabolic stability compared to control compounds (**Fig. S2B, C**). Additionally, we confirmed via SPR that our lead compounds AD-5584 and AD-8007 specifically bind ACSS2 and not the mitochondrial isoform hACSS1 (**Fig. S2D**).

### Evaluation of ACSS2 Inhibitors on Tumor Cell Growth and Lipid Content *In-Vitro*

Having identified compounds that bind to and inhibit ACSS2 *in vitro*, we sought to determine whether they had biological effects on brain tropic breast cancer cells. For this study, we utilize brain-trophic breast cancer cells, MDA-MB-231BR, which have been selected to preferentially metastasize to the brain and derived from parental breast cancer cells MDA-MB-231[39]. Our original candidate, AD-2441 was able to significantly reduce clonogenic survival in MDA-MB-231-BR cells 100 μM, similar to control VY-3-249 (**Fig.5A**). Following modifications to AD-2441, we tested the analogs AD-7346, AD-5584, AD-8007, AD-1363 at 100 μM where AD-5584, AD-8007, AD-1363 showed significant reduction in clonogenic survival (**Fig. 5B**). Since ACSS2 plays an integral role in generating acetyl-CoA from acetate, a substrate vital for lipid synthesis [21], we examined effect of novel ACCS2 inhibitors on lipid content. By quantifying lipid droplets, we found AD-8007 and AD-5584 compounds significantly reduced lipid droplet content in MDA-MB-231-BR cells when compared to the control treatment (**Fig.5C**). Since targeting ACSS2 blocked BCBM clonogenic cell survival, we tested whether our top candidates drugs AD-8007 and AD-5584 could increase cell death in MDA-MB-231-BR cells. Treatment of MDA-MB-231-BR cells with AD-8007 and AD-5584 significantly increased the percent of propidium iodide (PI) positive cell death compared to control and was comparable to effects seen with VY-3-135 **(Fig. 5D).** Thus, new ACSS2 inhibitors AD-5584 and AD-8007 block colony survival, reduce lipid content and induce cell death in BCBM cells.

**Figure 5:**
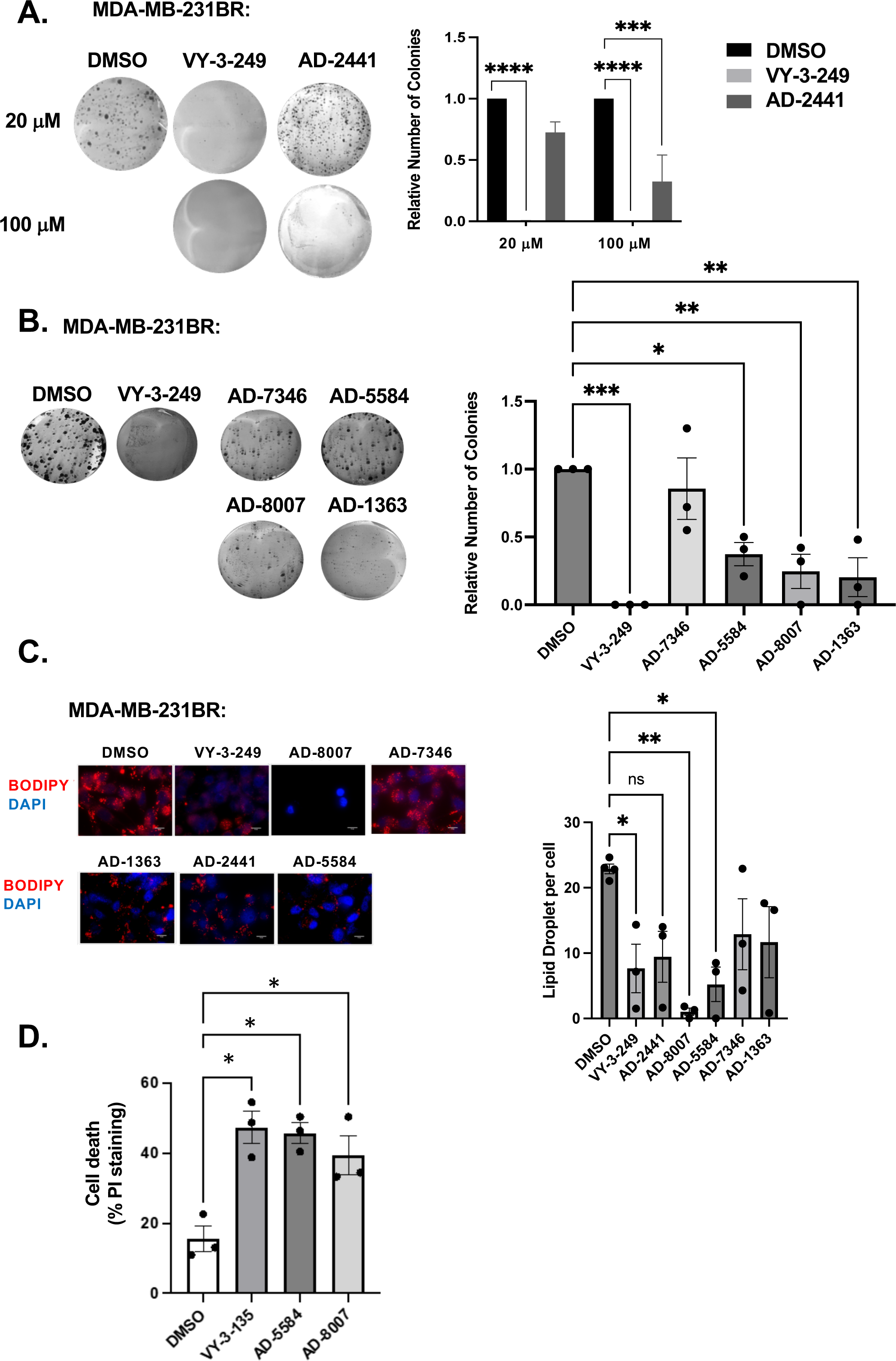
*In Vitro* effect of ACSS2 inhibitors on MDA-MB-231BR cells. (**A**) Representative images of MDA-MB-231BR cells treated with ACSS2 inhibitors for 48 hours at 100 μM and seeded in clonogenic cells survival assay stained with crystal violet, average colony formation quantified and presented as average from three independent experiments. Two-way ANOVA reported as mean ± SEM. ***p<0.001, ****p<0.0001. (**B**) Representative images of MDA-MB-231BR cells treated with ACSS2 inhibitors for 48 hours and seeded clonogenic cells survival assay stained with crystal violet, average colony formation quantified and presented as average from three independent experiments. One-way ANOVA reported as mean ± SEM. *p<0.05, **p<0.01, ***p<0.001. (**C**) Representative images of MDA-MB-231BR cells stained with BOPIDY following 48 hour treatment with ACSS2 at 100 μM and presented as average lipid droplet per cell from three independent experiments. One-way ANOVA reported as mean ± SEM. *p<0.05, **p<0.01, ***p<0.001. (**D**) Quantified graph of PI+ cells detected by flow cytometry analysis of MDA-MB-231BR cells treated with ACSS2 inhibitors for 48 hours. One-way ANOVA reported as mean ± SEM. *p<0.05, **p<0.01, ***p<0.001

### Evaluation of ACSS2 Inhibitors on BCBM Growth *Ex Vivo* and synergy with radiation

To test these compounds in a more physiologically relevant model, we employed our recently developed *ex vivo* tumor-brain slice model [40]. Treatment of *ex vivo* brain slices containing MDA-MB-231-BR cells with ACSS2 inhibitors AD-5584 and AD-8007 significantly reduced tumor growth from preformed tumors, suggesting induction of cell death compared to controls (**Fig. 6A**). Inhibition of tumor growth by novel ACSS2 inhibitors was similar to that detected by treatment of brain slices containing BCBM cells with VY-3-135 (**Fig. 6A**). Additionally, treatment of brain tissue, not containing tumors, with ACSS2 inhibitors AD-8007 and AD-5584 did not alter cell viability compared to positive control paraformaldehyde (**Fig. S3**). Thus, our novel ACSS2 inhibitors block tumor growth in the brain microenvironment while causing no toxicity to normal brain tissue.

**Figure 6:**
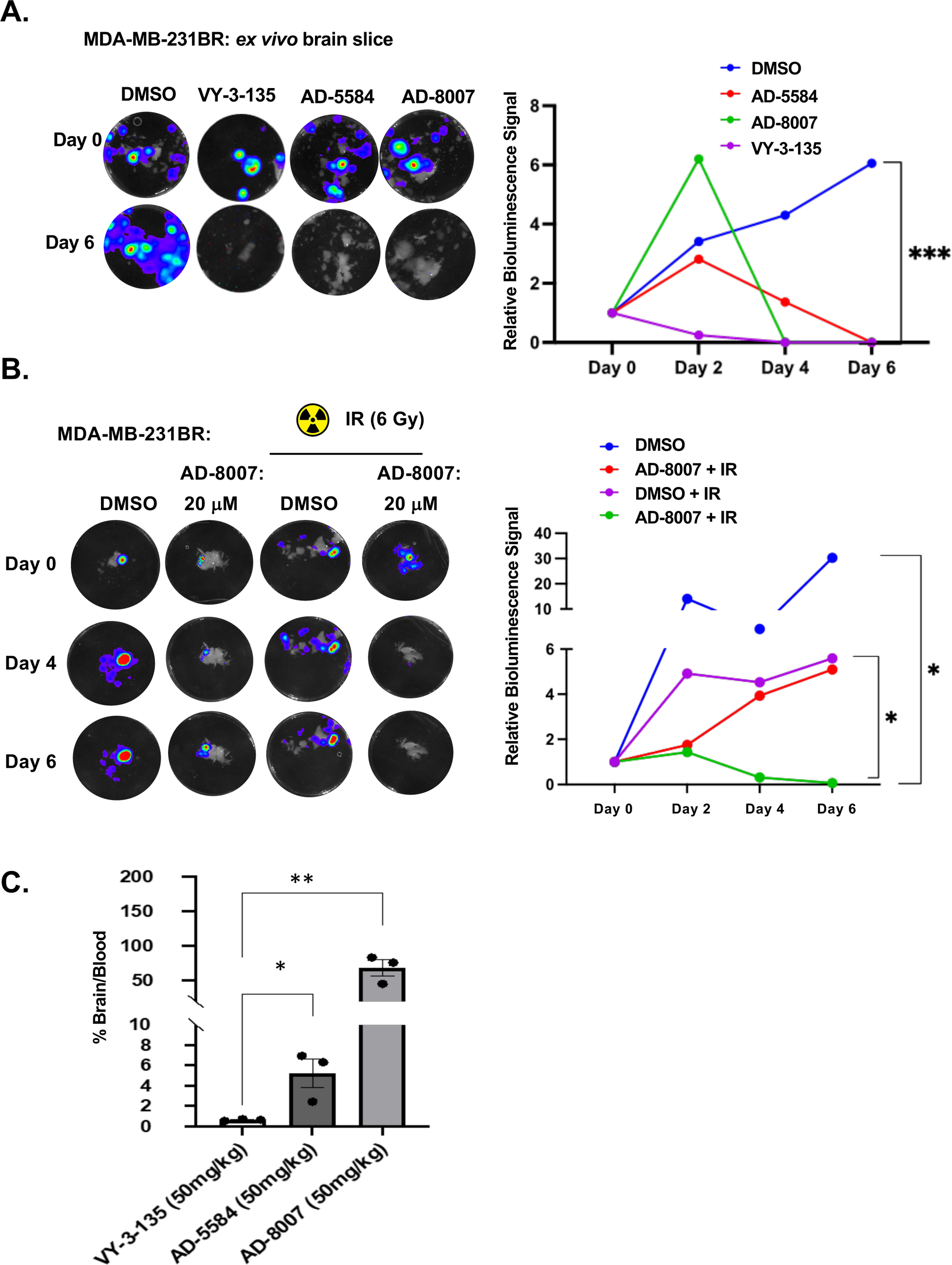
*Ex vivo* and BBB permeability of ACSS2 inhibitors. **(A)** Representative images of ex vivo tumor-brain slices were obtained from nu/nu mice injected with luciferase-tagged MDA-MB-231BR cells treated with ACSS2 inhibitors at 100 μM for 6 days. Quantified graph of relative bioluminescence signal at indicated day (n=3) Two-way ANOVA reported as mean ***p<0.001 **(B)** Representative images of *ex vivo* tumor-brain slices derived from nu/nu mice injected with luciferase tagged MDA-MB-231BR cells exposed to no irradiation (control) or one dose of 6Gy treated with ACSS2 inhibitors at 20 μM for 6 days (n=3). ANOVA reported as mean. *p<0.05. **(C)** Quantification of LC-MS peaks of ACSS2 inhibitor present in the brain over blood following intraperitoneal delivery of 50mg/kg of drug for 30 mins (AD-5584) or 1 hour (AD-8007, VY-3-135), blood extraction via intracardiac injection, perfusion, and brain retrieval. Student’s paired t-test reported as mean ± SEM. *p<0.05, **p<0.01.

Radiation is one of the first lines of treatment for patients with breast cancer brain metastasis [41]. We have previously shown that irradiation of brain slices containing MDA-MB-231BR leads to cytostatic effects *ex vivo* [40]; thus, we were interested in testing whether our new ACSS2 inhibitors can synergize with radiation in blocking BCBM growth. Treatment of preformed tumors with suboptimal dose of AD-8007 (20 μM) had some effect on BCBM cell growth *ex vivo* (**Fig. 6B**) and, as previously shown, treatment with 6 Gy radiation caused a cytostatic effect on BCBM growth (**Fig. 6B**). However, combination treatment of AD-8007 and radiation significantly blocked BCBM growth *ex vivo* (**Fig. 6B**). Thus, these data suggests that our novel ACSS2 inhibitors may be able to synergize with radiation to block BCBM growth and survival.

### Metabolic Stability and Blood-Brain Barrier Penetration Assessment of AD-5584 and AD-8007

We first evaluated the metabolic and plasma stability of AD-5584 and AD-8007 using human liver microsomes (HLMs). We included AD-3766 (not further assessed otherwise) as a control to validate our computational pipeline’s capacity to predict metabolic stability. Consistent with our predictions (**Fig. S2**), experimental validation indicated that both AD-5584 (T_1/2_ of 20min) and AD-8007 (T_1/2_ of >145min) demonstrated significantly higher metabolic stability than the control, AD-3766 (T_1/2_ of 0.9min) (**Table S2**). These findings suggest that AD-5584 and AD-8007 are promising foundations for further optimization, including potency and BBB permeability.

To evaluate the BBB permeability of AD-5584 and AD-8007, we used the human multidrug resistance protein 1 (MDR1)-transfected Madin-Darby canine kidney (MDCK) cell line (MDR1-MDCK) BBB assay [42]. This assay provides the apparent permeability coefficient (Papp), efflux ratio, and % recovery to assess compound permeation. AD-5584 and AD-8007 showed moderate permeability, with AD-8007 also displaying a low efflux ratio, indicating its potential to bypass P-gp substrate detection and cross the BBB (**Table S3**). These results align well with our computational predictions (**Sup Fig. 1C**).

Our *in vitro* and *ex vivo* results revealed the potency of AD-5584 and AD-8007 as novel ACSS2 inhibitors able to block growth, reduce lipid content, and induce cell death in BCBM cells. However, successfully translating these results to an *in vivo* context hinges on the compounds’ ability to cross the blood-brain barrier (BBB), reach their intended target, and remain metabolically stable. Although BBB permeability increases during brain metastasis progression due to the formation of a more permeable tumor-brain barrier (TBB)[43], leveraging therapeutics at the earliest possible stage while the BBB remains intact is a key clinical objective. Thus, optimal treatment candidates should demonstrate high BBB permeability *in vivo*. Our modeling predicted that AD-5584 and AD-8007 may be more brain-permeable compared to VY-3-135 (**Fig. S1C**). To test this *in vivo*, we injected mice with these compounds intraperitoneally and measured levels of drugs in plasma compared to brain homogenate. Utilizing LC-MS analysis, we found that our novel ACSS2 inhibitors AD-5584 and AD-8007 are detected at significantly higher levels in the brain compared to VY-3-135 at 50 mg/kg dose (**Fig. 6C**). Thus, we have successfully identified and validated AD-5584 and AD-8007 as novel ACSS2 inhibitors that reduce BCBM growth *in vitro* and *ex vivo*, reduce lipid content and induce cell death in MDA-MB-231BR cells. Importantly, both compounds demonstrated strong metabolic stability and the capacity to penetrate the blood-brain barrier *in vivo*.

## Discussion

Our study has identified and characterized novel ACSS2 inhibitors that can mitigate BCBM growth *in vitro* and *ex vivo* and are brain permeable *in vivo*. By applying a computational pipeline to screen and predict drug-like properties, we have identified and validated two potential compounds, AD-5584 and AD-8007, as specific inhibitors of human ACSS2. The computational pipeline effectively filtered potential ACSS2 binders from a pool of molecules. This *in silico* approach has leveraged unique computational techniques and software to predict ADME properties and other drug-like features. As *in silico* screening techniques have the potential to accelerate drug discovery and reduce costs, this study adds to the growing evidence supporting their utility.

Furthermore, our compounds of interest, AD-5584, and AD-8007, demonstrated improved drug-like characteristics and metabolic stability compared to the control compounds. This presents the promising potential for these compounds to withstand first-pass metabolism and reach their target site, essential attributes for drugs intended for systemic administration. However, it should be noted that the findings are predictive or *in vitro* validated and need further validation through *in vivo* metabolic stability assessments. In addition to identifying potential ACSS2 inhibitors, this study has also shed light on their mechanism of action. The identified compounds bind directly to ACSS2 and interfere with its role in lipid metabolism, as evidenced by the reduction in lipid droplet content within the MDA-MB-231BR cells. Given the integral role that ACSS2 plays in providing acetyl-CoA, a substrate vital for lipid synthesis, the potential of these inhibitors to interfere with lipid metabolism and, consequently, tumor growth is significant. Indeed, recent studies have shown that breast tumors in the brain must adapt to the low lipid availability in the brain by increasing *de novo* fatty acid synthesis and targeting the enzyme fatty acid synthase (FASN) can block breast cancer growth in the brain [44]. However, targeting FASN has proven problematic as many FASN inhibitors have failed to advance in the clinic due to largely unexpected *in vivo* toxicities [45]. Interestingly, the most potent ACSS2 inhibitors, AD-5584 and AD-8007, were found to induce cell death in MDA-MB-231BR cells but were not toxic in normal brain tissue, underscoring their potential as therapeutic candidates for BCBM. Future studies will explore the possible toxicities of these compounds.

In assessing the BBB penetration potential of these compounds, the study has also factored in the physiological context of drug delivery, particularly considering the crucial role of BBB permeability for drugs intended to act on brain targets. The BBB is a major hurdle for delivering many drugs to the brain, and it can become more permeable during brain metastasis progression. While BBB permeability can be leveraged during this stage of progression, for optimal treatment efficacy, drugs should be able to penetrate the BBB at the earliest stage possible. In this context, the BBB permeability of AD-5584 and AD-8007 holds promise. In addition, we show that AD-8007 can synergize with radiation treatment to block tumor growth *ex vivo*. Radiation treatment of BCBM often leads to cytostatic effects in patients [41] that we have shown can be modeled *ex vivo* [40] (**Fig. 6B**). Future studies will focus on evaluating tumor growth inhibition and induction of apoptosis *in vivo* to strengthen the potential of these compounds as effective and well-tolerated therapeutic candidates against BCBM as single agents and test synergy with radiation *in vivo*.

In conclusion, this study has identified brain-permeable ACSS2 inhibitors that can mitigate BCBM growth. The potential of the identified inhibitors, AD-5584 and AD-8007, their impact on lipid metabolism and induction of cell death, along with their high BBB permeability and metabolic stability, present encouraging avenues for further investigation and development as potential therapeutic candidates for BCBM. We aim to optimize our hit compounds further to reach a clinically relevant range, focusing on exceptionally metabolically stable AD-5584 and AD-8007 and further developing these new drugs for treating patients with cancer brain metastasis.

## Methods

### Cell culture

Triple-negative brain trophic cells MDA-MB-231BR are a kind gift from Dr. Patricia Steeg (Center for Cancer Research, National Cancer Institute) and were cultured in DMEM supplemented with 10% fetal bovine serum (FBS), 5% 10,000 Units/mL Penicillin-10,000 μg/mL Streptomycin, and 5% 200 mM L-Glutamine. ACSS2 inhibitors were dissolved in 100% ultra-pure DMSO.

### Clonogenic survival assay

To assess clonogenic survival, 1 x 10^5^ cells were seeded into a 6 well plate until 70-80% confluency and treated for 48 hours with analogs at 100 uM (unless otherwise stated in the figure legend). After 48 hours, cells were trypsinized and counted using hemocytomer, and 1×10^4^ cells per treatment were plated into a 6 well plate with fresh culture medium and allowed to grow for 10 days. Following 10-day incubation, cells were washed twice with PBS, stained with crystal violet for 30 minutes, and washed with dH_2_O twice. Colonies >50mircons were counted. For crystal violet staining, 0.5% crystal violet was prepared in a 1:1 methanol-water solution.

### BODIPY staining of cells

Cells were treated with 5uM BODIPY 493/503 in PBS for 15 min, washed 2x with 1xPBS, fixed using 4% paraformaldehyde for 30 min at RT in the dark and washed in 1xPBS prior to mounting and imaging on EVOS FL (Life Technologies) using Texas Red filter.

### Flow Cytometry

Cells were prepared according to manufacturer protocol (BD Pharmingen FITC Annexin V Apoptosis Detection Kit). Briefly, cells were trypsinized (0.25% Trypsin), counted, washed twice with 1xPBS, and resuspended in 100uL 1X binding buffer incubated with 5ul Propidium Iodine staining solution for 15 minutes in the dark at room temperature. Following incubation, the volume was brought up to 500uL of 1X binding buffer. Tubes were then analyzed using a Guava easyCyte flow cytometer. All data were collected and analyzed using a Guava EasyCyte Plus system and CytoSoft (version 5.3) software (Millipore). Data are gated and expressed relative to the appropriate unstained and single-stained controls.

### *Ex vivo* brain slice model

Nu/Nu athymic 6-8 week old mice Charles River Laboratories (Wilmington, MA, USA) were immobilized using the Just for Mice Stereotaxic Frame (Harvard Apparatus, Holliston, MA, USA) and injected intracranially with 5uL of 100,000 cells/uL solution of luciferase tagged MDA-MB-231BR cells. Tumor growth was monitored via bioluminescence imaging on the IVIS system (Perkin Elmer, Waltman, MA, USA). Organotypic hippocampal cultures were prepared as described previously (ref). Briefly, adult mice (6-8 week) or mice after 12 days following intracranial injection were decapitated and their brains rapidly removed into ice-cold (4°C) sucrose-aCSF composed of the following (in mM): 280 sucrose, 5 KCl, 2 MgCl_2_, 1 CaCl_2_, 20 glucose, 10 HEPES, 5 Na^+^-ascorbate, 3 thiourea, 2 Na^+^-pyruvate; pH=7.3. Brains were blocked with a sharp scalpel and sliced into 250 µm sections using a McIlwain-type tissue chopper (Vibrotome Inc.). Four to six slices were placed onto each 0.4 µm Millicell tissue culture insert (Millipore) in six-well plates, 1 ml of medium containing the following: Neurobasal medium A (Gibco), 2% B27 supplement, 1% N2 supplement (Gibco), 1% glutamine (Invitrogen), 0.5% glucose, 10 U/ml penicillin, and 100 ug/ml streptomycin (Invitrogen), placed underneath each insert. The media was changed every 2 days following imaging. Tumor growth was monitored via bioluminescence imaging on the IVIS 200 system (Perkin Elmer), and results were analyzed using Living Image software (Caliper Life Sciences, Waltham, MA, USA). For the MTS assay, individual brain slices were transferred to a 96-well plate and subjected to Promega CellTiter 96^®^ Aqueous One Solution (Cat: G3582) mixed in a 1:5 ratio with culture media and treated as previously described. Tissues were incubated at 37℃, 5% CO_2_ for 4 hours, and absorbance at 490nm was measured with a Tecan Spark Microplate reader.

### BBB permeablity *in vivo*

Human ACSS2 inhibitors were prepared in 10 mg/mL saline solution. BalbC 6-8 week-old mice were weighed and injected with 50 mg/kg or 100 mg/kg of drug saline solution via intraperitoneal injection. After 30 minutes or 60 mins, mice treated with AD-5584 or AD-8007 and VY-3-135 were placed in isoflurane, and blood was extracted and perfused via intracardiac injection. Following perfusion, mice were decapitated, and their brains were rapidly removed into ice-cold PBS.

For analysis of blood samples, 200 µl of blood was transferred and allowed to clot for 15 minutes at room temperature. Samples were precipitated at 16000xg for 1 minute, and 50 µl of serum was transferred. To serum, 200 µl of methanol was added, and samples were centrifuged at 16000xg for 5 minutes. The supernatant was analyzed by LCMS. For analysis of brain samples, brains were homogenized via bead beating using four 2.3 mm stainless steel beads in 1 ml of MeOH and 0.5 ml of H_2_O. After centrifugation for 15 min at 3000xg, the supernatant was analyzed by LCMS. LCMS was performed on an Acquity I-Class UPLC system coupled to a Synapt G2Si HDMS mass spectrometer in positive ion mode with a heated electrospray ionization (ESI) source in a Z-spray configuration. LC separation was performed on a Waters Acquity UPLC BEH 1.7 μm 2.1×50mm column equipped with a Vanguard guard column, using a 0.6 mL/min solvent flow of A/B 95/5 to 15/85 in 4 minutes, followed by washing and reconditioning the column. Eluent A is 0.1% v/v formic acid in water, and B is 0.1% v/v formic acid in acetonitrile. Conditions on the mass spectrometer were as follows: capillary voltage 0.5 kV, sampling cone 40, source offset 80, source 120°C, desolvation temperature 250°C, cone gas 0, desolvation gas 1000 L/hr and nebulizer 6.5 bar. The analyzer was operated in resolution mode. Low energy was collected between 100 and 1500 Da at 0.2 sec scan time. MSe data was collected using a ramp trap collision energy 20-40 V. Masses were extracted from the TOF MS TICs using an abs width of 0.05 Da. Data was analyzed using Waters MassLynx and Waters Unifi. Calibration curves of authentic standards were used for quantification.

### Overproduction and purification of human ACSS2

Overproduction of ACSS2 was achieved using a prokaryotic expression system. Briefly, the plasmids containing the C- and N-terminally His-tagged human ACSS2 DNA were transformed into BL21 (DE3) RIL competent cells (Agilent Technologies, Wilmington, DE) and were expressed in auto-inducing media ZYP-5052 overnight at 15°C with shaking at 225 rpm. The bacterial expressions were then spun down, the supernatant discarded, and the pellets resuspended in 50 mM Tris HCl pH8.0, 800 mM NaCl, 5 mM Imidazole, 2 mM MgCl_2_, 10% Glycerol, 0.1 mg/mL Lysozyme, 1 mM Protease inhibitor (PMSF). After the cells were lysed via sonication, the sample was subjected to ultracentrifugation, and the clarified lysate was applied to a 5 mL Talon cobalt resin affinity column (Clonetech Laboratories, Mountain View, CA). The bound protein was washed with 300 mL wash buffer 1 (20 mM Tris HCl pH8.0, 800 mM NaCl, 15 mM Imidazole, 5 mM MgCl_2_, 10% Glycerol, 0.1% CHAPS) and 300 mL wash buffer 2 (20 mM Tris HCl pH8.0, 400 mM NaCl, 18 mM Imidazole, 5 mM MgCl_2_, 10% Glycerol) prior elution with 20 mM Tris HCl pH8.0, 400 mM NaCl, 300 mM Imidazole, 2 mM MgCl_2_, 5% Glycerol. Eluted ACSS2 was then dialyzed overnight into 20 mM Tris HCl pH8.0, 150 mM NaCl, 5% Glycerol, concentrated and applied to a Superdex S-200 (16/600) using the same buffer without Glycerol. ACSS2 fractions were pooled, concentrated to 3 mg/mL, aliquoted, and stored at −80°C. hACSS1 was obtained from from a commercial vendor (www.mybiosource.com).

### SPR characterization

All binding assays were performed on a ProteOn XPR36 SPR Protein Interaction Array System (Bio-Rad Laboratories, Hercules, CA, USA). The instrument temperature was set at 25 °C for all kinetic analyses. ProteOn GLH sensor chips were preconditioned with two short pulses each (10 s) of 50 mM NaOH, 100 mM HCl, and 0.5% sodium dodecyl sulfide. Then, the system was equilibrated with running buffer (1x PBS pH 7.4, 3% DMSO, and 0.005% polysorbate 20). The surface of a GLH sensor chip was activated with a 1:100 dilution of a 1:1 mixture of 1-ethyl-3-(3-dimethylaminopropyl) carbodiimide hydrochloride (0.2 M) and sulfo-N-hydroxysuccinimide (0.05 M). Immediately after chip activation, the human ACSS2 or hACSS1 proteins were prepared at a concentration of 10 μg/mL in 10 mM sodium acetate, pH 5.5, and injected across ligand flow channels for 5 min at a flow rate of 30 µL/min. Then, after unreacted protein had been washed out, excess active ester groups on the sensor surface were capped by a 5 min injection of 1M ethanolamine HCl (pH 8.0) at a flow rate of 5 μL/min. A reference surface was similarly created by immobilizing a nonspecific protein (IgG b12 anti-HIV-1 gp120; was obtained through the NIH AIDS Reagent Program, Division of AIDS, NIAID, NIH: Anti-HIV-1 gp120 Monoclonal (IgG1 b12) from Dr. Dennis Burton and Carlos Barbas) and was used as a background to correct nonspecific binding. Serial dilutions of ACSS2 inhibitors or a single concentration at 25 μM of AD-5584, AD-8007, and VY-3-249 for hACSS1 binding were then prepared in the running buffer and injected at a flow rate of 100 µL/min for a 50 s association phase, followed by up to a 5 min dissociation phase using the “one-shot kinetics” capability of the ProteOn instrument. Data were analyzed using the ProteOn Manager Software version 3.0 (Bio-Rad). The responses from the reference flow cell were subtracted to account for the nonspecific binding and injection artifacts. Experimental data were fitted to a simple 1:1 binding model. Experiments were performed in triplicate to detect kinetic and equilibrium dissociation constants (K_D_).

### Fluorescence polarization-based ACSS2 biochemical assay (ATP to AMP conversion)

ACSS2 enzyme activity was measured using the TranScreener AMP^2^/GMP^2^ Assay Kit – FP Readout assay (BellBrook Labs). The assay was performed in white, opaque, 96-well plates. Compounds diluted in 100% DMSO were used starting at 150 nM with a 1:3 dilution, and ACSS2 was used at 100 nM. ACSS2 was used in assay buffer (30 mM HEPES, pH 7.4, 140 mM NaCl, 2 mM MgCl_2_, 5 mM sodium acetate, 2 mM DTT, 0.05% CHAPS). Substrate mix was added, followed by a 60-minute incubation. Final substrate concentrations were 5 mM acetate, 50 μM ATP, and 5 μM CoA. After incubation, conjugated AMP antibody and AMP tracer were added according to the methods described by BellBrook Labs. After 30 minutes, the FP signal was measured using a Tecan Spark multimode microplate reader. In the analysis, data were normalized to represent the percentage inhibition of ATPase activity. A value of 100% inhibition corresponded to the counts observed in the absence of ACSS2, whereas 0% inhibition was aligned with the counts from the complete reaction, including a DMSO control.

### Pharmacophore-based Shape Screen and AD-2441 Analog Identification (HT Screening)

The reference Quinoxaline molecule [15] was drawn and prepared in VIDA 5.0.4.0 (OpenEye, Cadence Molecular Sciences, Santa Fe, NM. http://www.eyesopen.com) and then exported to Szybki 2.6.0.1 (OpenEye, Cadence Molecular Sciences, Santa Fe, NM. http://www.eyesopen.com) for in solution minimization to be used as the lowest possible conformer. The ChemBridge diversity library was downloaded from their database website (https://chembridge.com) and prepared by Szybki 2.6.0.1 (OpenEye, Cadence Molecular Sciences, Santa Fe, NM. http://www.eyesopen.com). Then, about 200 conformers were generated for each molecule in the library using Omega Szybki 4.2.2.1 [46](OpenEye, Cadence Molecular Sciences, Santa Fe, NM. http://www.eyesopen.com). ROCS Szybki 3.6.0.1 [47](OpenEye, Cadence Molecular Sciences, Santa Fe, NM. http://www.eyesopen.com) was then used to build the 3D query for the reference Quinoxaline molecule, and then we screened the prepared ChemBridge library conformers against the query. The step was repeated twice to validate the hits and select the top 5000 hits. We then used StarDrop V7.3 (Optibrium Ltd. Cambridge, UK) to predict the drug-like properties (using the CNS penetration module) for preselection. To facilitate and improve confidence for hit selection, we continued with a structure-based docking approach, including predicting binding affinity (using the HYdrogen Bond and DEhydration Energies (HYDE) function)[48] in SeeSAR 12.1 and the homology model (swissmodel.expasy.org) of ACSS2 based on the crystal structure of ACSS2 from Salmonella enterica (PDB: 5JRH) which was prepared using Flare, version 5 (Cresset®, Litlington, Cambridgeshire, UK, http://www.cresset-group.com/flare/) to select the top 30 molecules for experimental evaluation.

After we evaluated the first 30 hits and identified AD-2441, we used two approaches for analog identification. The first approach included a simple Tanimoto coefficient cutoff of >0.9 within the ChemBridge.com library with subsequent SeeSAR 12.1 HYDE binding affinity evaluation and StarDrop V7.3 for drug-like properties filtering. The second approach included a ligand-focused SAR approach using a field-based search within Forge V10.6 (Cresset®, Litlington, Cambridgeshire, UK) and the Diversity library from Chembridge, followed by SeeSAR 12.1 HYDE binding affinity evaluation and StarDrop V7.3 for drug-like properties filtering.

### Docking calculations for ACSS2 inhibitors (Figure preparation)

The ACSS2 inhibitors were prepared, and then energy minimized using Flare version 5 (Cresset®, Litlington, Cambridgeshire, UK, http://www.cresset-group.com/flare/) with a root mean squared (RMS) gradient cutoff of 0.2 kcal/mol/A and 10000 iterations. The homology model of ACSS2 based on the crystal structure of ACSS2 from Salmonella enterica (PDB: 5JRH) was prepared using Flare, version 5 (Cresset®, Litlington, Cambridgeshire, UK, http://www.cresset-group.com/flare/) to allow protonation at pH 7.0 and removal of residue gaps. Docking calculations were performed using DiffDock [49] (github.com/gcorso/DiffDock), a diffusion generative model, using a blind docking approach (grid box over the entire protein). During the docking procedure, we used 100 inference steps and 300 samples per complex with a batch size of 12. Based on the DiffDock confidence score and SMINA score, the best 3 complexes were chosen. While DiffDock provides exceptional accuracy for blind docking, we noticed difficulties in the quality of the final poses. Therefore, we used Flare, version 5, to energy minimize the best complex form DiffDock, using the accurate XED force field minimization algorithm with a gradient cutoff of 0.050 kcal/mol/A and 10,000 iterations. In order to validate the quality of our homology model and DiffDock docking approach, we docked Adenosine-5’-propylphosphate (extracted from PDB: 1PG4) and compared the pose to the crystal structure (PDB: 1PG4).

### Statistical Analysis

All results shown are results of at least three independent experiments and are shown as averages and presented as mean ± s.e (or SEM if stated). P-values were calculated using a Student’s two-tailed test (* represents p-value ≤ 0.05 or **p-value ≤ 0.01 or as marked in figure legend). Statistical analysis of the growth rate of mice was performed using ANOVA. *p-value < 0.05.

## Acknowledgments

This work was supported by NIH-NCI grant UO1CA244303 (to MJR), PA Breast Cancer Coalition Award (to MJR), and Coulter-Drexel Translational Research Award (to AD & MJR)

## Author contributions

EE performed the *in vitro, ex vivo,* and *in vivo* work. LC tested AD-2441 and early analogs *in vitro.* RY, JM, and AT helped with the experimental work. JB performed LCMS on plasma and brain tissue. AD performed computation analysis, SPR, biochemistry, and docking studies. NLS helped with radiation experiments. EE, LC, AD, and MJR participated in the design, and data analysis of the study. AAR, SC, AD, and MJR participated in the conception of the study. All co-authors reviewed the final manuscript version.

## Supplementary Figures

**Figure S1:**
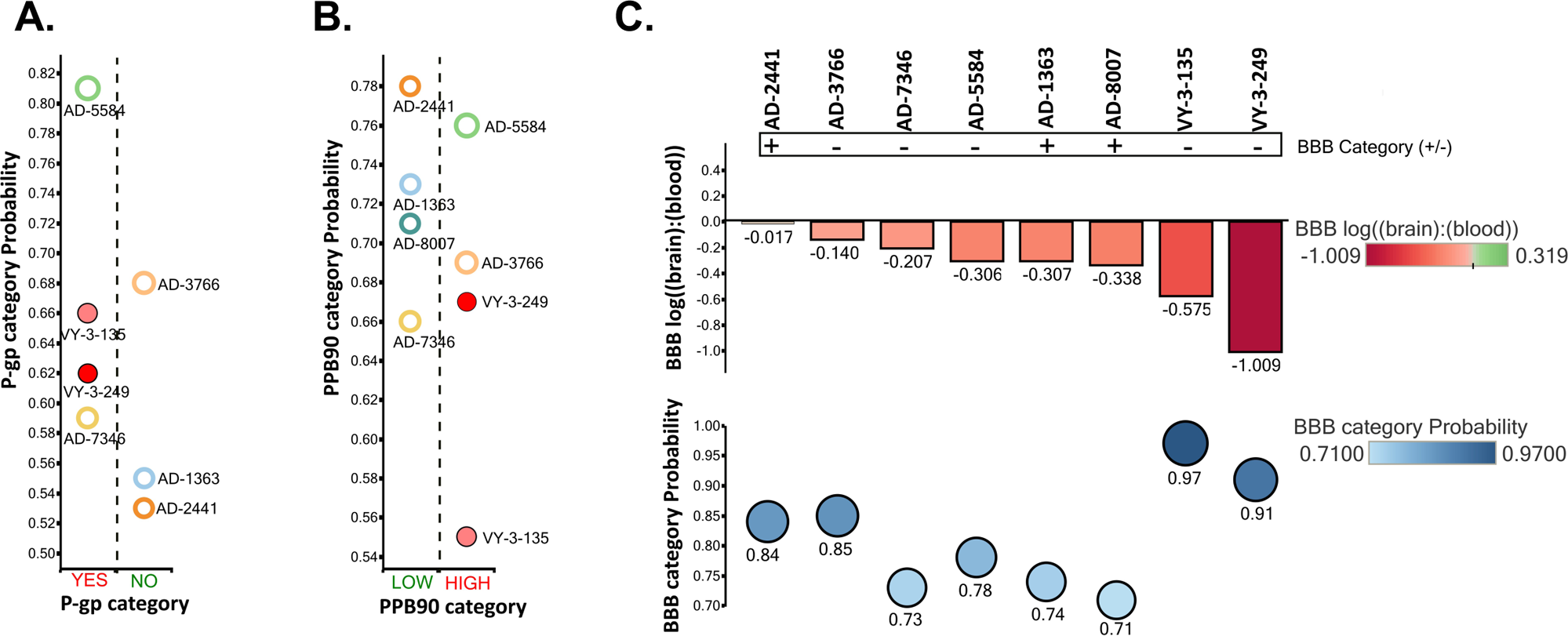
Computational prediction of drug-like properties of AD-2441 and its analogs. **(A)** P-glycoprotein (Pgp) category and predicted probability of AD-2441 and analogs**. (B)** Predicted plasma protein binding (90% threshold, PPB90) category and predicted probability. (**C)** Predicted blood-brain barrier (BBB) distribution/category and category-probability of AD-2441 and analogs.

**Figure S2:**
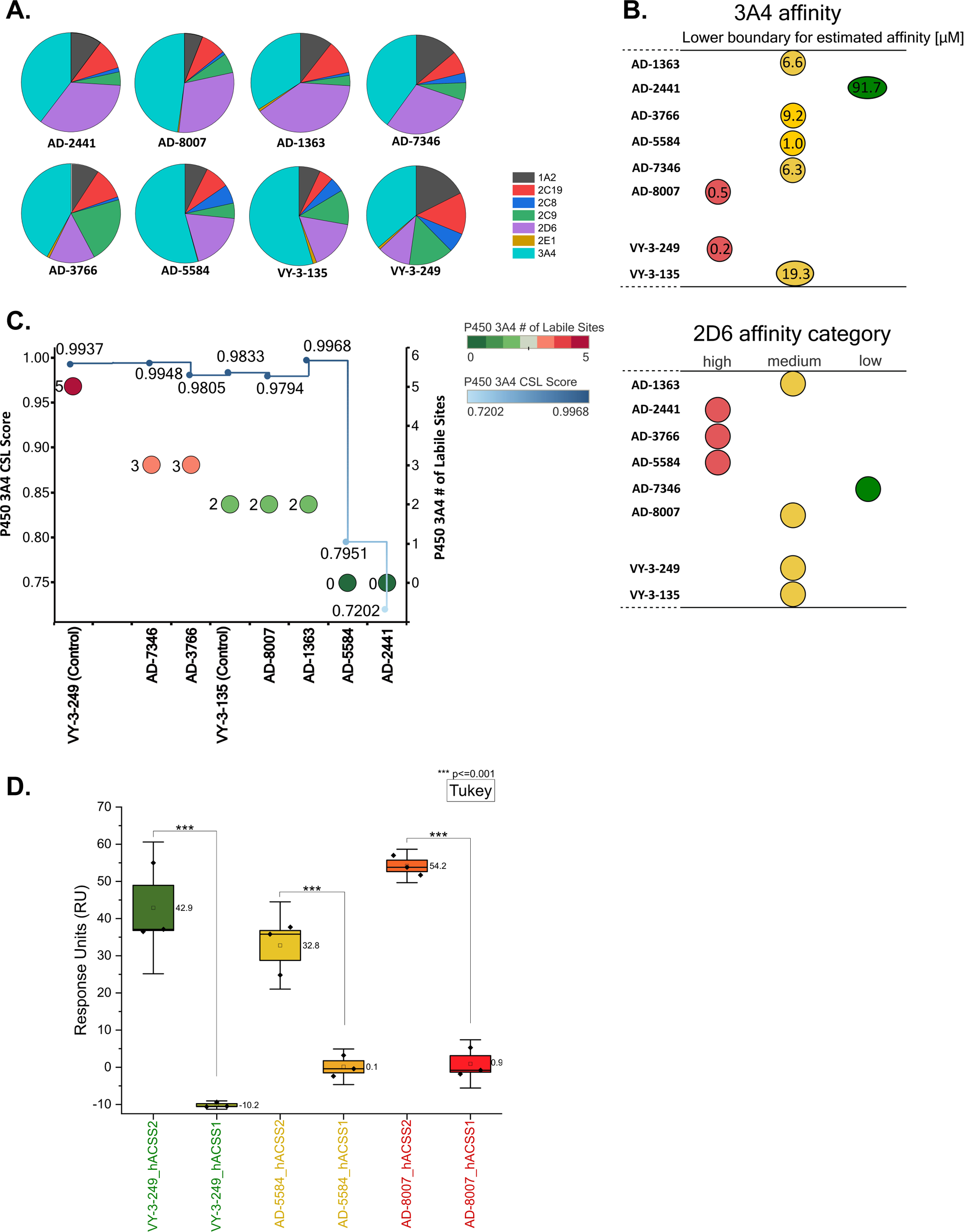
Computational prediction of metabolic stability of AD-2441 and its analogs. **(A)** Prediction of the major metabolizing CYP isoforms for AD-2441 and analogs. The majority of compounds are predicted to be metabolized by the 3A4 isoform, except AD-1363, which is predicted to be metabolized by the 2D6 isoform. (**B**) A lower boundary for predicted 3A4 affinity of AD-2441 and analogs using the hydrogen bond and dehydration scoring function (HYDE) implemented in SeeSAR12.1. 2D6 affinity category of AD-2441 and analogs as predicted by the Stardrop P450 module. (**C**) Overall composite site lability (CSL) score and number of labile sites (for metabolism) for AD-2441 and analogs. A lower CSL score indicates a more stable molecule. The prediction was achieved using the StarDrop (version 7) P450 module. **(D)** SPR-derived Response Units (RU) at 25 μM injection of VY-3-249, AD-5584, and AD-8007 to immobilized human ACSS2 or human ACSS1. Experiments were performed in triplicate (n=3), and statistical significance was performed using paired comparison and means compared using the Tukey method. Boxblots display the mean (line in the box and value to the right), box size represents SEM, and whiskers the confidence interval at α of 90%.

**Figure S3:**
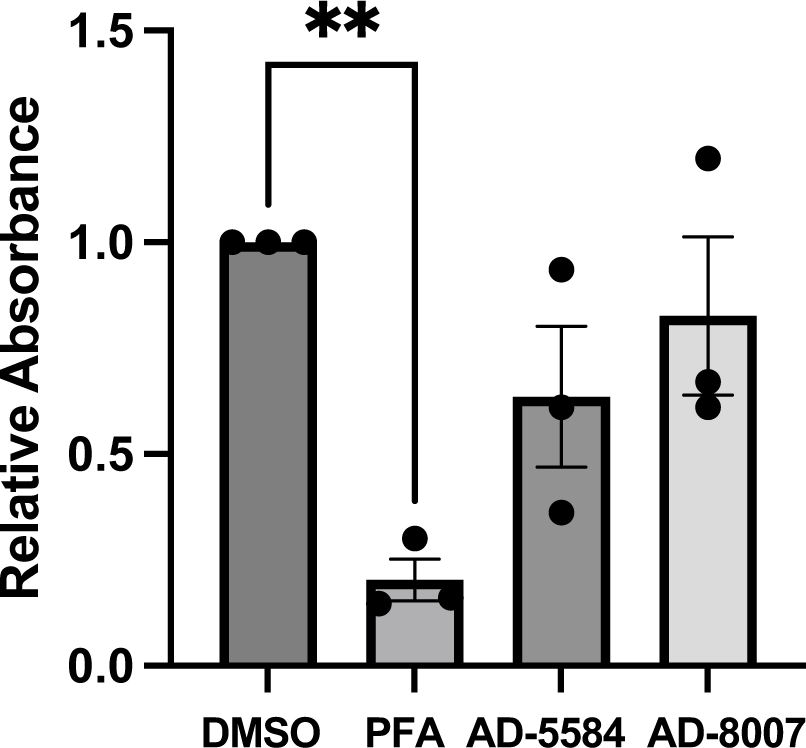
AD-5584 and AD-8007 are not cytotoxic to the brain tissue. *Ex vivo* brain slices were treated as in 6A (without tumor) with either DMSO or 100 μM of AD-8007 or AD-5584, collected on day 6, and analyzed for cell viability (MTS assay). As a positive control, slices were treated with paraformaldehyde (PFA), rendering the brain slices non-viable (n = 3). Student’s t-test reported as mean ± SEM. * p < 0.05.

**Table S1:**
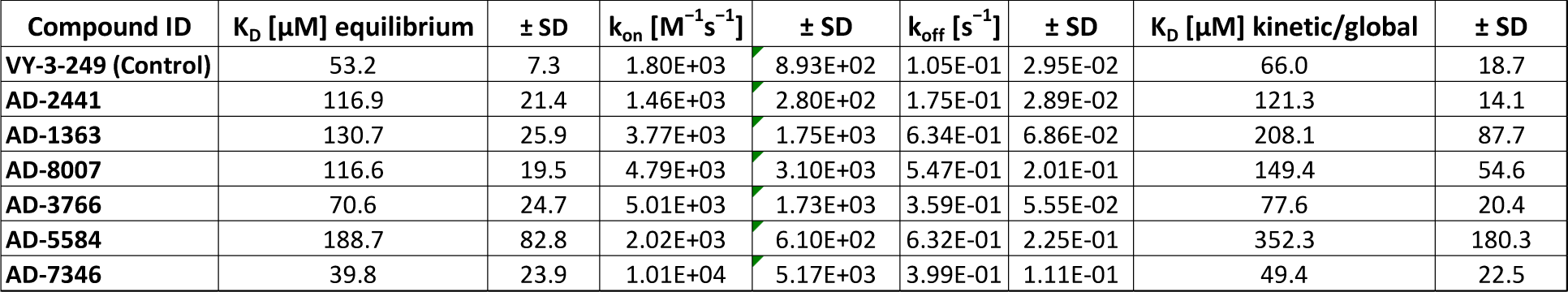
Kinetic and equilibrium parameters for AD-2441 and analogs binding to immobilized ACSS2. Equilibrium Dissociation Constant (K_D_) derived from a Langmuir isotherm equilibrium fit or a global fit. Values are mean ± standard deviation (SD) with n=3.

| Compound ID | $K_D$ [μM] equilibrium | ± SD | $k_{on}$ [$M^{-1}s^{-1}$] | ± SD | $k_{off}$ [$s^{-1}$] | ± SD | $K_D$ [μM] kinetic/global | ± SD |
| --- | --- | --- | --- | --- | --- | --- | --- | --- |
| VY-3-249 (Control) | 53.2 | 7.3 | 1.80E+03 | 8.93E+02 | 1.05E-01 | 2.95E-02 | 66.0 | 18.7 |
| AD-2441 | 116.9 | 21.4 | 1.46E+03 | 2.80E+02 | 1.75E-01 | 2.89E-02 | 121.3 | 14.1 |
| AD-1363 | 130.7 | 25.9 | 3.77E+03 | 1.75E+03 | 6.34E-01 | 6.86E-02 | 208.1 | 87.7 |
| AD-8007 | 116.6 | 19.5 | 4.79E+03 | 3.10E+03 | 5.47E-01 | 2.01E-01 | 149.4 | 54.6 |
| AD-3766 | 70.6 | 24.7 | 5.01E+03 | 1.73E+03 | 3.59E-01 | 5.55E-02 | 77.6 | 20.4 |
| AD-5584 | 188.7 | 82.8 | 2.02E+03 | 6.10E+02 | 6.32E-01 | 2.25E-01 | 352.3 | 180.3 |
| AD-7346 | 39.8 | 23.9 | 1.01E+04 | 5.17E+03 | 3.99E-01 | 1.11E-01 | 49.4 | 22.5 |

**Table S2:** Metabolic Stability of selected ACSS2 inhibitors in human liver microsomes and Plasma stability (WuXi AppTec Co., Ltd.). Testosterone, Diclofenac, Propafenone, and Propantheline Bromide were used as controls.

| Compound ID | Human Liver Microsome Stability |  |  |  |  | Human Plasma Stability |  |
| --- | --- | --- | --- | --- | --- | --- | --- |
|  | R<sup>2</sup> | T<sub>1/2</sub> (min) | CL<sub>int</sub>(mic)(μL/min/mg) | CL<sub>int</sub>(liver)(mL/min/kg) | Remaining(T=60min) | Remaining(NCF=60min) | T<sub>1/2</sub> (min) |
| AD-8007 | 0.9824 | >145 | <9.6 | <8.6 | 0.77227 | 0.95633 | >289.1 |
| AD-3766 | 1 | 0.914 | 1516.099 | 1364.4891 | 0.0005 | 0.96386 | >289.1 |
| AD-5584 | 0.983 | 19.565 | 70.842 | 63.7578 | 0.12368 | 0.83493 | >289.1 |
| Testosterone | 0.9884 | 15.519 | 89.309 | 80.3781 | 0.06959 | 0.89758 |  |
| Diclofenac | 0.9939 | 5.293 | 261.875 | 235.6875 | 0.00044 | 0.87629 |  |
| Propafenone | 0.9182 | 7.919 | 175.015 | 157.5135 | 0.00396 | 0.93177 |  |
| Propantheline Bromide |  |  |  |  |  |  | 18.6 |

**Table S3:** *In vitro* permeability assessment of AD-8007 and AD-5584 in MDR1-MDCK1 cells and appropriate controls (Creative Bioarray, Shirley, USA)

Bi-directional permeability across MDR1-MDCK I cell monolayer - Mimiking the BBB
| Compound ID | Mean P<sub>app</sub> (10<sup>-6</sup> cm/s) |  | Efflux Ratio | Mean %Solution Recovery |  | Note |
| --- | --- | --- | --- | --- | --- | --- |
|  | A to B | B to A |  | A to B | B to A |  |
| Nadolol | 0.257 | ND | ND | 99.1 | ND | Low permeability marker |
| Metoprolol | 12.9 | ND | ND | 94.2 | ND | High permeability marker |
| Digoxin | 0.0595 | 7.74 | 130 | 84.3 | 100.4 | P-gp substrate |
| AD-8007 | 2.81 | 25.2 | 8.96 | 81.3 | 97.0 | AD-8007 has a low efflux ratio and potential to cross the BBB |
| AD-5584 | 2.11 | 44.4 | 21.0 | 94.9 | 111.6 | --- |

ND means not determined

